# Feasibility Study Utilizing NanoString’s Digital Spatial Profiling (DSP) Technology for Characterizing the Immune Microenvironment in Barrett’s Esophagus FFPE Tissues

**DOI:** 10.1101/2023.11.02.565001

**Authors:** Qurat-ul Ain, Nicola Frei, Amir M Khoshiwal, Pim Stougie, Robert Odze, Lorenzo Ferri, Lucas C Duits, Jacques Bergman, Matthew D Stachler

## Abstract

To date, characterization of the Barrett’s esophagus (BE) immune microenvironment in patients with known progression status to determine how the microenvironment may influence BE progression to esophageal adenocarcinoma (EAC) has been understudied, hindering both the biological understanding of progression and the development of novel diagnostics and therapies. Therefore, this study’s aim was to determine if highly multiplex interrogation of the immune microenvironment can be performed on endoscopic formalin-fixed, paraffin-embedded (FFPE) samples utilizing the Nanostring GeoMx digital spatial profiling (GeoMx DSP) platform. We performed spatial proteomic analysis of 49 proteins expressed in the microenvironment and epithelial cells of histologically identical FFPE endoscopic biopsies from patients with non-dysplastic BE (NDBE) who later progressed to high-grade dysplasia (HGD) or EAC (N=7) or from patients who after at least 5 years follow up did not (N=8). In addition, we performed RNA analysis of 1,812 cancer related transcripts on a series of three endoscopic mucosal resections containing regions of normal tissue, BE, dysplasia (DYS), and EAC. Our primary goal was to determine feasibility of this approach and begin to identify the types of specific immune cell populations that may mediate the progression of pre-neoplastic BE to EAC. Spatial proteomic and transcriptomic profiling with GeoMx DSP showed reasonable quality metrics and detected expected differences between epithelium and stroma. Several proteins were found to have increased expression within non-dysplastic BE biopsies from progressors compared to non-progressors, suggesting further studies on the BE microenvironment are warranted.

**Summary:** New biological insights into the stepwise development and progression of esophageal adenocarcinoma (EAC) from Barrett’s esophagus (BE) are imperative to develop tailored approaches for early detection and optimal clinical management of the disease. This study aimed to determine the feasibility to spatially profile stromal and immunologic properties that accompany malignant transformation of BE to EAC in formalin-fixed, paraffin-embedded (FFPE) tissues. NanoString’s Digital Spatial Profiling (DSP) technology can detect and quantify protein and RNA transcripts in a highly multiplexed manner with spatial resolution, within specific regions of interest on FFPE tissue. Here, we performed a pilot study using the Nanostring GeoMx DSP, for measurement of protein and ribonucleic acid (RNA) expression on a series of FFPE slides from endoscopic biopsies and endoscopic mucosal resections (EMR) of BE. We compare a small series of biopsies of non-dysplastic BE (NDBE) from patients who progressed to more advanced disease to patients with NBDE who did not progress and then perform RNA profiling on EMRs with a range of histologic diagnoses.

## Introduction

Barrett’s esophagus (BE) is the premalignant stage of esophageal adenocarcinoma (EAC) [1–3]. EAC remains one of the deadliest forms of gastro-intestinal cancer with a mortality rate exceeding 90% for advanced disease [4, 5]. While the exact process of EAC formation is not fully understood, it is thought to develop sequentially along the metaplasia-dysplasia - adenocarcinoma sequence with progression attributed to a series of genomic and epigenetic events that ultimately allow for clonal selection of BE cells [6–10]. Prior studies on the molecular mechanisms underlying EAC and its stepwise progression from BE have primarily focused on tumor cell-intrinsic features, such as genomic alterations. These studies have made significant progress in our understanding of BE progression [11,12]. However, other extrinsic factors likely also play a role in the progression process. Together with genomic alterations, there is accumulating evidence that the inflammatory microenvironment (iME) plays a role in the progression process by contributing to inhibiting apoptosis, enabling immune evasion, and promoting proliferation, angiogenesis, invasion, and metastasis [13, 14]. Studies in the microenvironment of BE have revealed involvement of lymphoid cells (i.e., T cells, B cells, NK cells) and myeloid-derived cells (i.e., macrophages, neutrophils, eosinophils), as well as their secreted cytokines and chemokines[15, 16]. So far, however, the immune contexture (type, density, and location, as well as phenotypic and functional profile of immune cells) has not been extensively investigated to understand how it may be involved in the progression of BE. A deeper understanding of how BE/EAC-iME interactions contribute to tumorigenesis may direct the development of future diagnostic and therapeutic strategies in high-risk patients to inform clinical decision-making.

While traditional immunohistochemistry (IHC) techniques allow for the spatial profiling of cells, this is often lost when samples are analyzed using broader profiling techniques such as RNA sequencing (either bulk or single cell approaches). To this end, the local cellular proportions, cellular heterogeneity, and deeper spatial distribution are often lacking in tumor microenvironment characterization studies [17]. Combining spatial, cellular composition, and cell state information can aid in identifying micro-niches within the iME that may allow for deeper insights or associations to be identified that can be lost in bulk approaches such as rarer immune cells within the epithelium versus within the stroma. The determination of the immune makeup within the iME lays the foundation for addressing how the immunological composition and status (activated/suppressed) may dictate BE progression or response to therapy. Therefore, to address this need, proper tissue sampling along with technologies that retain spatial information while providing hi-plex capabilities are both required.

In this study, we utilized the GeoMx digital spatial profiler (GeoMx DSP) technology to evaluate its effectiveness to investigate the iME within BE and during progression. The GeoMx DSP is a system for proteomics and transcriptomics analyses in a spatially resolved manner developed by NanoString Technologies (Seattle, WA) [18]. The GeoMx DSP applies several advanced technologies, including high-throughput readout, programmable digital micromirror device for precise area selection, and microfluidic sampling, creating an innovative tool for discovery, translational research, and clinical uses [19].

Our goal was to conduct a pilot study to quantify the abundance of tumor and immune-related transcriptional and protein markers in separate epithelial and stromal compartments predominately focused on non-dysplastic Barrett’s esophagus (NDBE) but also including samples with dysplasia and early EAC. We utilized the Cancer Transcriptome Atlas (CTA) for high-plex analysis of messenger ribonucleic acids (mRNAs) and a panel of 49 antibodies for protein analysis. We apply this method to study formalin fixed paraffin embedded (FFPE) biospecimens selected from a large cohort of NDBE endoscopic biopsies as well as endoscopic mucosal resections. Further we aimed to demonstrate the ability of this tool for gastrointestinal (GI) research through investigating the molecular underpinnings and immunomodulatory pathways that underlie BE progression to EAC.

## Materials and Methods

### Samples selection

FFPE blocks of endoscopic biopsies of varying age were selected from a previously described cohort and FFPE blocks of endoscopic mucosal resections (EMR) retrieved from pathology archives of the University of San Francisco or the McGill University Hospital system [20]. Fifteen biopsy blocks with a histologic diagnosis of NDBE from 7 patients who after at least two years progressed to high-grade dysplasia (HGD) or EAC and 8 patients who after a minimum of 5 years of follow-up did not progress were utilized for protein analysis. Three EMR blocks with a range of histologic diagnoses from NDBE to early EAC were utilized for mRNA analysis. The study was performed after IRB approval (UCSF# 19-27460).

### Tissue Processing and hematoxylin and eosin (H&E) Staining

Five-micron sections were taken from each block. The first and last section was stained with hematoxylin and eosin (H&E) following the procedure previously described [21] and reviewed by trained GI pathologists (MDS) to confirm histologic diagnosis in each stained section. Supplementary Figure 1 shows representative hematoxylin and eosin images of the endoscopic biopsies (Supplementary Figure 1A) and EMRs (Supplementary Figure 1B).

### Sample Preparation and GeoMx DSP Profiling

The study was performed as part of the Nanostring Technology Access Program (TAP) and all slide processing was performed within the Nanostring facility (Seattle, WA). Unstained 5 µm thickness tissue sections were shipped to Nanostring where they were processed according to previously published protocols for protein and RNA analysis [22].

#### Spatial Transcriptomics Analysis

To analyze mRNA transcripts in spatial context we utilized the human Cancer Transcriptome Atlas (CTA) which profiles 1812 transcripts selected for their relevance in immunology and cancer. Briefly, slides were baked for 2 hours at 65°C for paraffin removal and subsequently loaded onto a Leica BOND RX (Leica Biosystems, Wetzlar, Germany) for tissue rehydration in EtOH and ddH_2_O, heat-induced epitope retrieval (ER2 for 20 minutes at 100°C) and proteinase K treatment (1.0 μg/ml for 15 minutes at 37°C). The tissue sections were then hybridized with the CTA probes overnight at 37°C. Following 2X 5min stringent washes (1:1 4x SSC buffer & formamide), the slides were blocked and then incubated with morphology marker antibodies to guide region of interest (ROI) selection: PanCK (488 channel, NBP2-33200AF488, Novus Biologicals), CD68 (594 channel, sc-20060AF594, Santa Cruz), and CD45 (647 channel, NBP2-34528AF647, Novus Biologicals). Syto83 (532 channel, S11364, Invitrogen) was used as a nuclear stain. Once the staining was completed, the slide was loaded on to GeoMx DSP instrument where they were scanned to produce an individual digital fluorescent image based on the morphology markers described above (Supplementary Figure 1C and 1D) as described in Nanostring protocol [22, 23] to enable morphology marker guided ROI selection. After ROI selection by a trained GI pathologist (MDS), the ROIs were further sub-segmented into areas of interest (AOI) based on PanCK staining (epithelium vs stroma) then UV light was directed by the GeoMx DSP at each AOI releasing the RNA-ID containing oligonucleotide tags from the CTA probes for collection into a unique well for each AOI. For library preparation, Illumina i5 and i7 dual indexing primers were added to the oligonucleotide tags through PCR to uniquely index each AOI. AMPure XP beads (Beckman Coulter) were used for PCR purification. Library concentration as measured using a Qubit fluorometer (Thermo Fisher Scientific) and quality was assessed using a Bioanalyzer (Agilent). Sequencing was performed on an Illumina NextSeq 550 and fastq files were processed into gene count data for each AOI using the GeoMx NGS Pipeline.

#### Spatial Proteomics Analysis

For protein analysis, unstained sections were interrogated with 49 antibodies focused predominately around immune cell types, immune activation, and immune modulation markers (Supplementary Table 1). Slides for protein analysis were treated similarly for paraffin removal, and tissue re-hydration using standard laboratory protocol [22,23]. However, antigen retrieval was done with citrate buffer, at high temperature and pressure. After washing in TBS-T, slides were blocked for 1 hour before morphology marker (same as CTA) addition. GeoMx Human IO antibodies (Immune Cell Profiling, IO Drug Targets, Immune Activation Status, Immune Cell Typing, Pan Tumor modules) were added for an overnight incubation at 4°C. After another TBS-T wash, slides were post-fixed in 4% PFA for 30 min at RT. Following an additional wash, slides were stained with ROI selection markers and underwent a similar process for ROI/AOI selection as described above. Post AOI selection, the oligo tags were cleaved from the panel antibodies and were hybridized with Probe R/U and Plex-Set Reporter Tags, before being counted on an nCounter MAX/FLEX system. RCC files were uploaded to the GeoMx for data analysis.

### Strategies for ROI Selection & Segmentation in Corresponding AOIs

Using the morphology markers and paired H&E slide, 6 individual regions of interest (ROIs) per slide were selected to encompass both the BE epithelium as well as the surrounding stroma/lamina propria of approximately ∼3 BE gland profiles in size. Areas which contained squamous epithelium or gastric type glands were avoided.

Using the Pan-Keratin morphology marker, each ROI was sub segmented into Pan-Keratin positive (PanCK+) AOIs for epithelium and Pan-Keratin negative (PankCK-) AOIs for stroma/microenvironment characterization as described by Van, T.M. and Blank, C.U [17], which were each analyzed separately (Supplementary Figure 1E and 1F). Anti-CD45 and anti-CD68 antibodies were utilized to help select representative areas. Once each ROI was selected and segmented into AOIs, slides were processed as described above.

### Data Quality Control of ROI/AOI and Data Normalization

After quantification of probes, data from the protein and CTA analysis was uploaded to the Nanostring GeoMx for QC, normalization, and analysis.

#### Protein QC and normalization

For the protein data, several different methods of normalization were compared to determine the most suitable. These included 3 housekeeping (HK) proteins (GAPDH, Histone H3 & Ribosomal protein S6), 3 immunoglobulin G (IgG) controls including (Ms IgG1, Ms IgG2a & Rb IgG), number of nuclei, and overall surface area. IgG and HK normalization account for both differences in ROI size and cellularity [24].

#### CTA QC and normalization

CTA data was first checked for successful sequencing. Second, the 5 probes for each CTA target were filtered for any outliers before consolidation into a single count value per sample per target. Counts were normalized using the third quartile (Q3) normalization method [24].

### Statistical analysis

After normalization, all statistical analysis and visualization was performed utilizing the built in GeoMx analysis software. Normalized log2 counts were used for the statistical analyses. For analyzing the expression correlation, the Pearson correlation coefficient (R coefficient) was reported for paired groups; R > 0 indicates a positive correlation, while R < 0 suggests a negative correlation. The expression difference between two groups was determined using linear mix modeling, while the one-way analysis of variance test was utilized for comparison across multiple groups. Normalized log2 counts were further zero-centralized for generating the heatmap. The Euclidian method was used to calculate the individual distance, and the average hierarchical clustering method was used to determine the expression clusters.

## Results

### Feasibility, normalization, and QC

For protein analysis, multiple methods of normalization were first tested and compared by correlation and standard deviation to determine the optimum approach. Analysis suggested normalization utilizing the housekeeping genes showed a strong correlation and was a good normalization strategy for the protein data (Figure 1) and is what was utilized to analyze the data.

**Figure 1:**
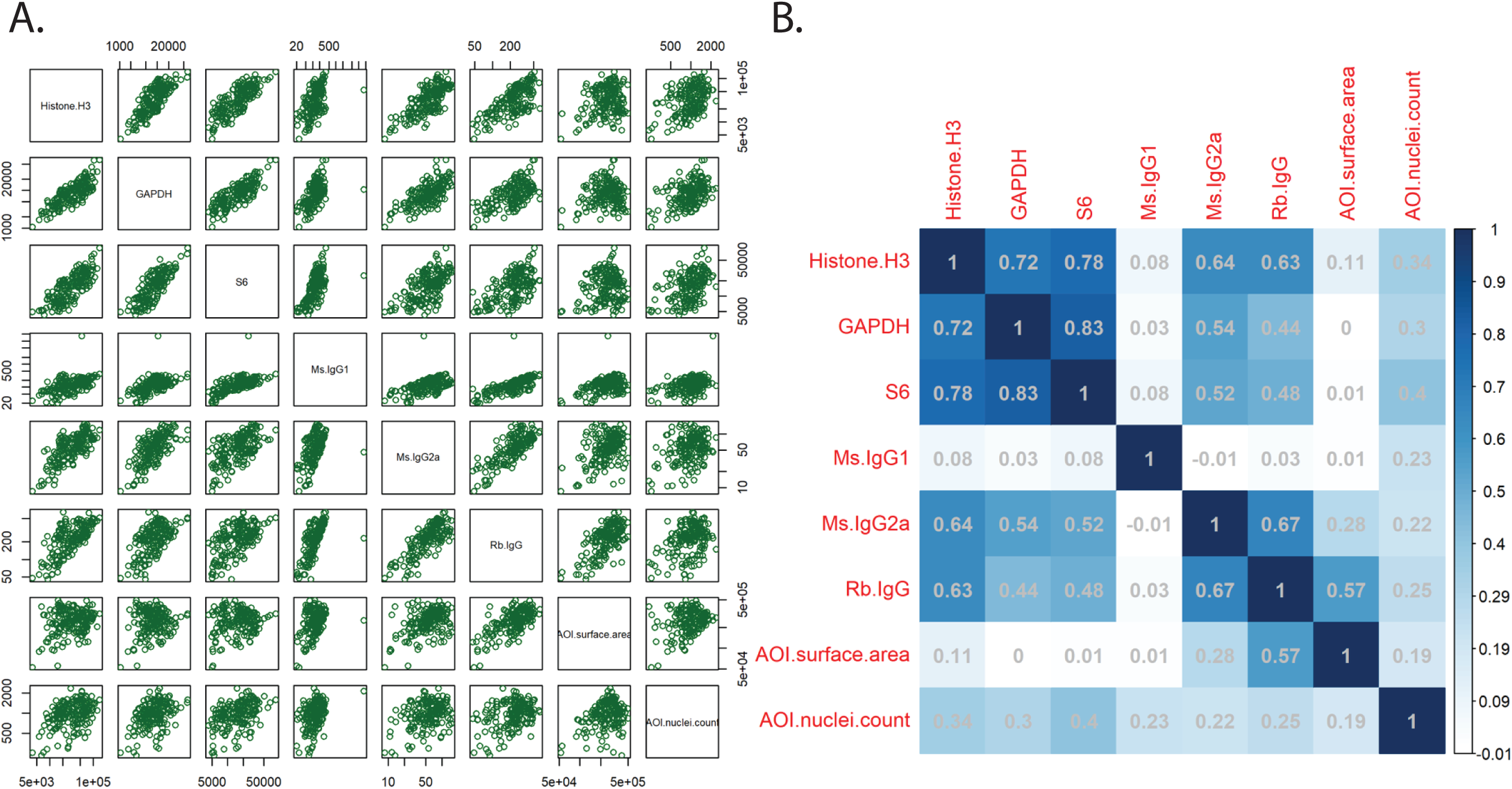
Normalization and QC of protein data. Normalized AOI count plots (A) and Pearson correlation coefficients (B) of the various normalization strategies, including housekeeping genes (Histone H3, GAPDH, ribosomal protein S6), IgG controls (mouse and rabbit), AOI surface area, and AOI nuclei count.

For RNA analysis, counts from each AOI were analyzed and no sample was considered under-sequenced for further analysis/processing. After normalization, 36/1,812 (2%) RNA targets were considered below the limit of quantification (LOQ). Either these RNA targeted are not expressed in the analyzed samples or the probes for that target failed. 368/1,812 (20%) targets were detected over the LOQ but in less than 20% of AOIs, potentially showing differential tissue expression or limited successful detection. The rest of the probes, 1,408/1,812 (78%), were detected above the LOQ in greater than 20% of the AOIs (Figure 2).

**Figure 2:**
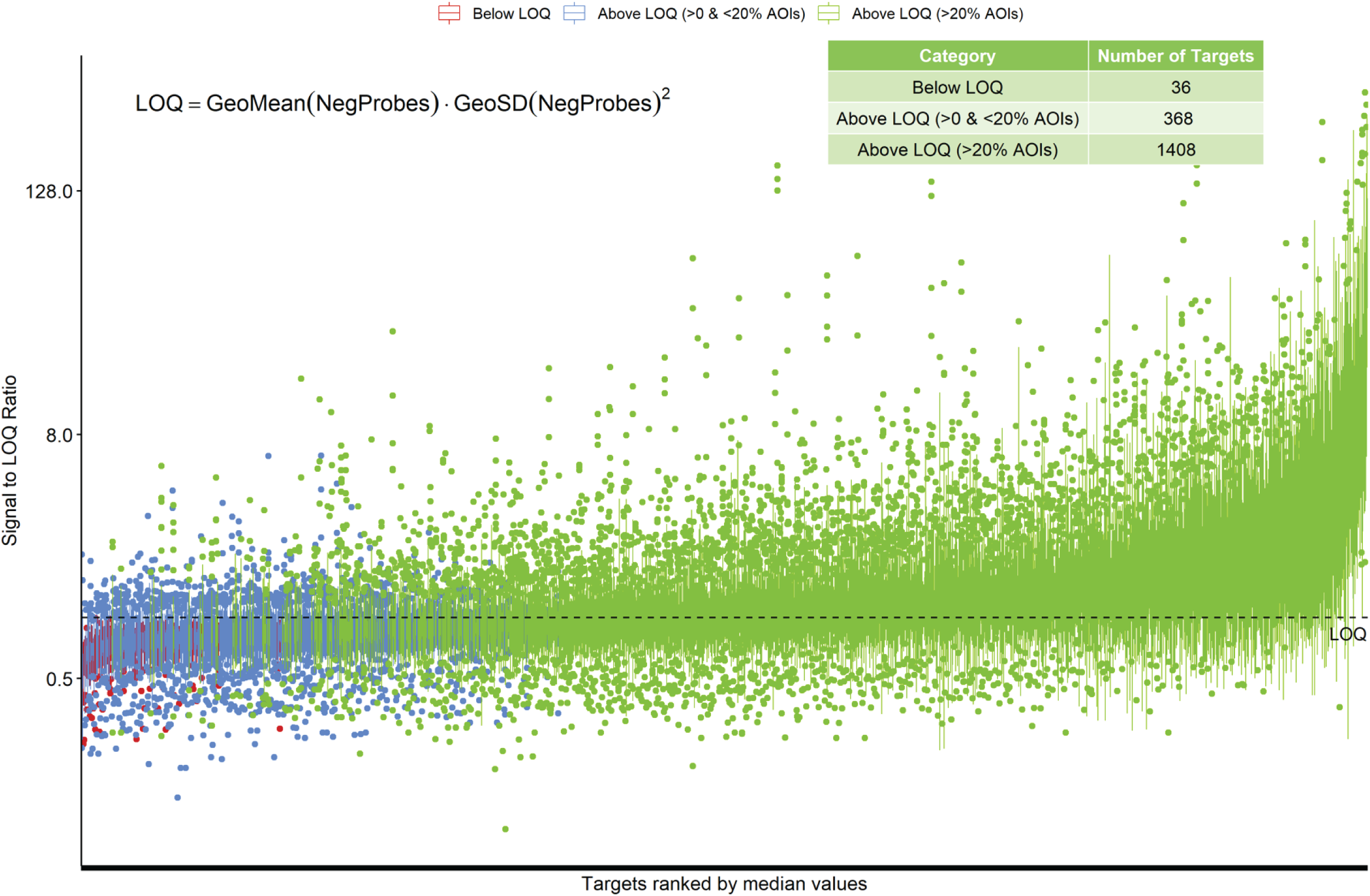
Normalization and QC of RNA data. Individual counts were normalized against the 75^th^ percentile of signal from their own AOI (Q3 normalization) and the limit of quantification (LOQ) was calculated. The normalized signal to LOQ was plotted for each RNA target. Red=below LOQ, blue=above LOQ in less than 20% of AOIs, green=above LOQ in greater than 20% of AOIs.

To ensure masking and AOI selection separated the epithelium and stromal areas appropriately, the expression of pan-cytokeratin (panCK) for protein and Krt19 for RNA was compared in the two components. As expected, both were highly overrepresented in the epithelial compartment (Figure 3A (Protein) 3B (RNA)).

**Figure 3:**
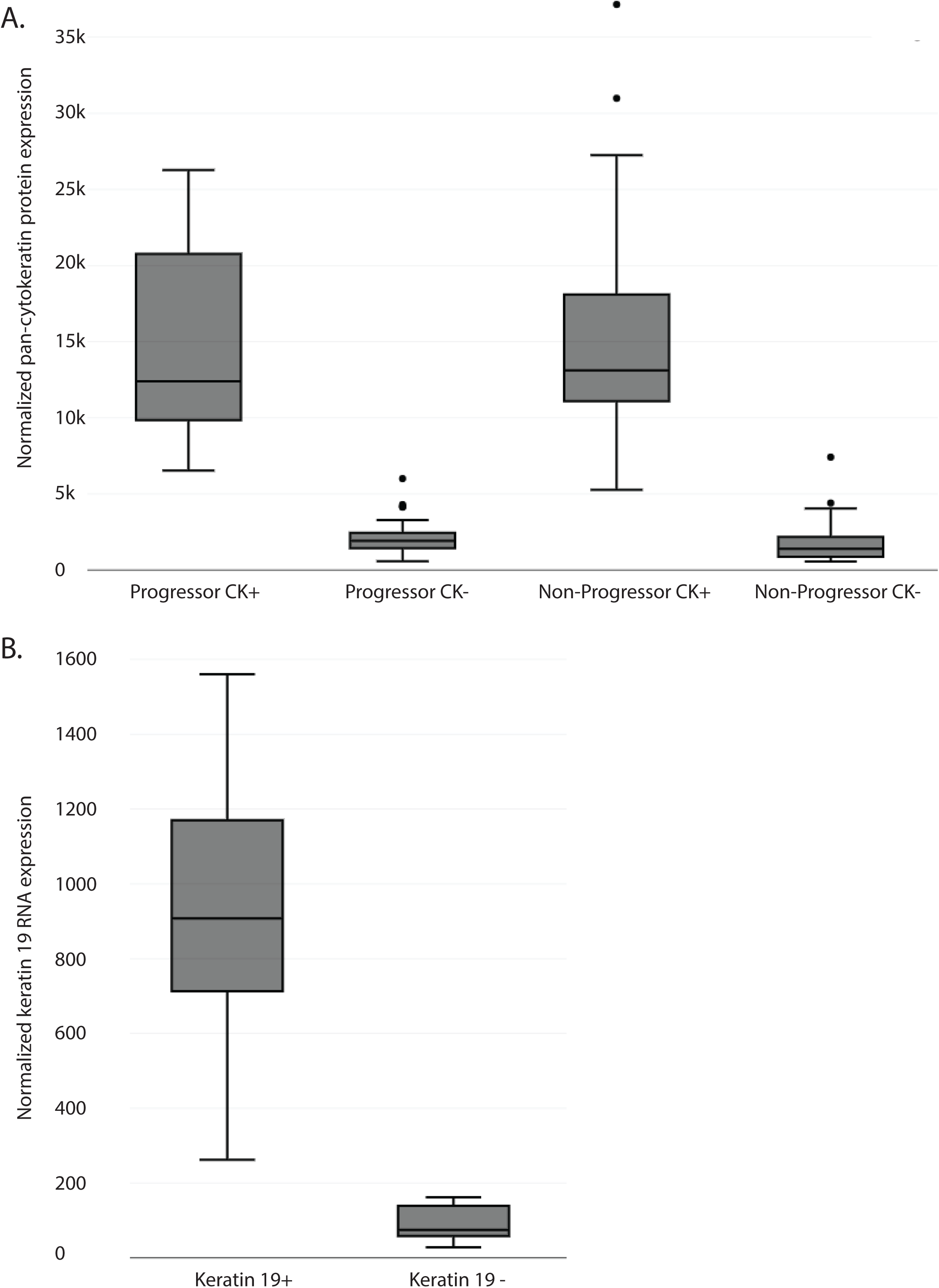
Segmentation of ROIs into Pan-CK+ and Pan-CK-AOIs. Slides were stained with the immunofluorescent morphology marker Pan-CK and used to segment selected ROIs into Pan-CK+ AOIs (Barrett’s epithelium) and Pan-CK-AOIs (surrounding stroma). Box plots show the normalized pan-CK protein expression (A) and keratin 19 RNA expression (B) of the different AOIs. Protein quantification was separated into progressing and non-progressing patients.

t-Distributed Stochastic Neighbor Embedding (tSNE) plots of the protein data revealed, as expected, a segregation of AOIs by type with the epithelial AOIs showing in a tight cluster (Supplementary Figure 2A). Additionally, Unsupervised clustering of all protein AOIs segregated the epithelial and stromal AOIs (Supplementary Figure 2B). Similarly, for the RNA analysis, when looking at differential gene expression between Pan-CK+ and Pan-CK-AOIs, the most upregulated genes in the epithelial compartment included keratins and EPCAM while multiple collagens, immune related transcripts, and VCAM1/PECAM1 were over-represented in the stromal compartment (Supplementary Figure 3).

### Differential protein expression in FFPE samples of non-dysplastic Barrett’s esophagus from progressing and non-progression patients

For protein analysis we isolated 42 ROIs from 7 patients who progressed and 48 ROIs from 8 patients who did not progress. While tSNE and unsupervised clustering was able to separate the epithelial AOIs from the stromal AOIs, it was unable to cleanly separate progressors from non-progressors. However, differential gene expression suggested several differences between progressors and non-progressors with several proteins being overexpressed in the progression samples (Figure 4). Within the epithelial compartment, several immunoregulatory proteins were upregulated in the progressors compared to the non-progressors, including CD45RO, HLA-DR, CTLA4, among others. Similar to the epithelial compartment, several immunomodulatory proteins were also found to be over-represented in the stroma of progressors compared to non-progressors, including PD-1, PD-L1, and CTLA4. While not completely specific, in addition to the immunomodulatory proteins, protein markers for several immune cells were also suggested to be over-represented in progressors compared to non-progressors. These included NK cells (CD56, GZMB), granulocytes/neutrophils (CD66b), macrophages (CD68), and CD4 T-cells (CD4) among others. In addition to the immune markers, SMA a protein associated with myofibroblasts and fibronectin a protein produced by a variety of cells including cancer associated fibroblasts showed increased expression within the stromal compartment of progressors compared to non-progressors.

**Figure 4:**
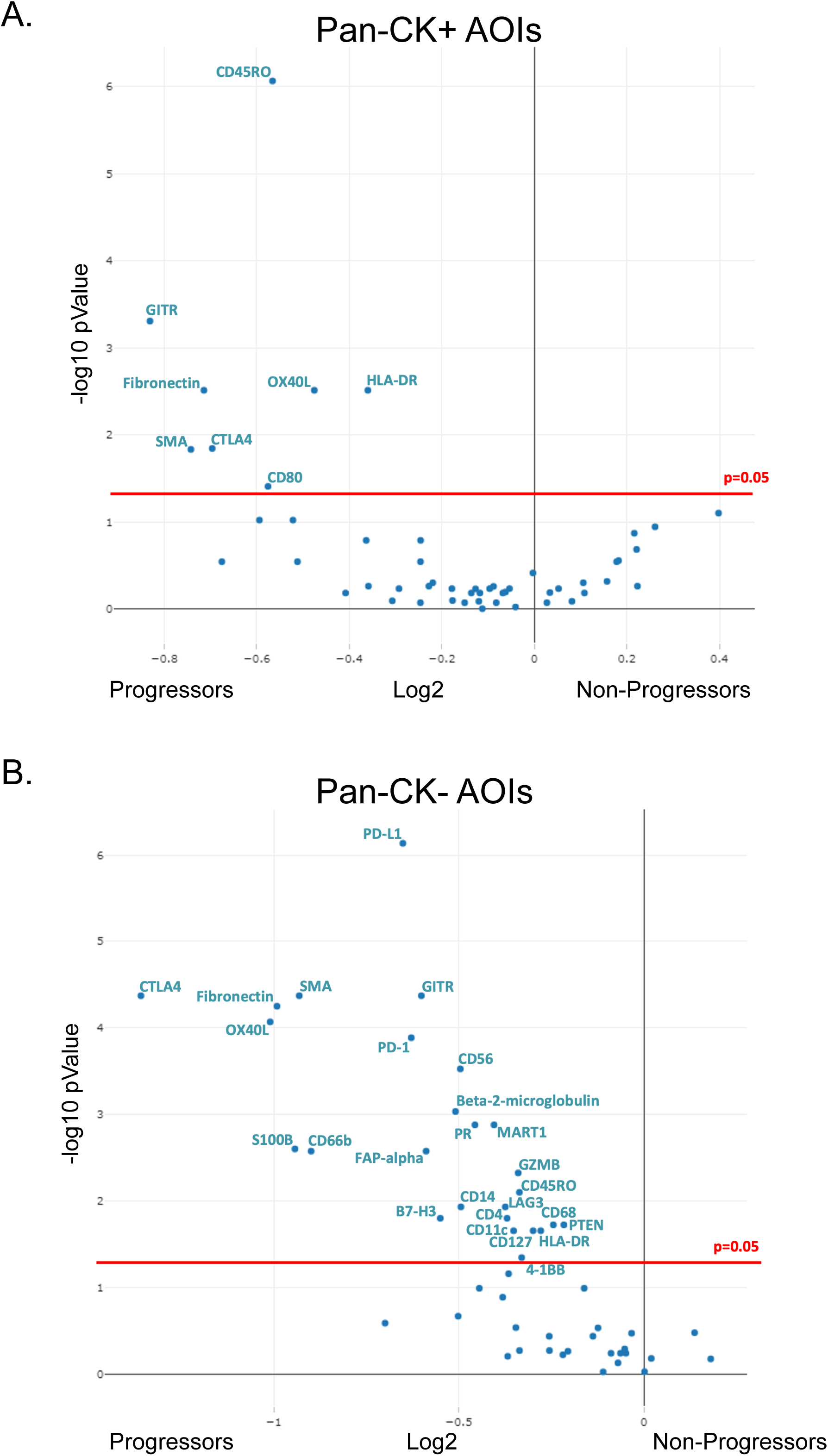
Differential protein expression in progressing and non-progressing BE patients. Volcano plots showing differential protein expression after Benjamini-Hochberg correction for multiple testing in pan-CK+ AOIs (A) and pan-CK-AOIs (B).

### Feasibility of spatial RNA detection in FFPE Barrett’s esophagus samples

To determine how the DSP platform performs in analyzing RNA from endoscopic specimens, four endoscopic mucosal resections with a variety of histologic diagnoses and the CTA was utilized. The CTA interrogates over 1,800 RNA transcripts covering multiple cancer and immune related pathways. Similar to the protein studies, ROIs of approximately similar size were selected using IF and a paired hematoxylin and eosin stained slide and then sub-segmented into AOIs using PanK. A total of 24 ROIs were selected (NDBE n=8, LGD/HGD n=8, and EAC n=8). While the number of samples was extremely small, when comparing the NDBE to the dysplastic epithelial AOIs, several transcripts were upregulated in NDBE compared to dysplastic AOIs including CD68, TMEM45B, and CCL28 while ILF3, MIF, DNMT1, and NPM1 were upregulated in the dysplastic AOIs (Figure 5A). When comparing the dysplastic to the EAC epithelial AOIs, multiple genes including CCND1(commonly amplified in EAC), BCL2L1 (apoptosis regulator), and multiple TNF superfamily members were upregulated in the EAC AOIs (Figure 5B).

**Figure 5:**
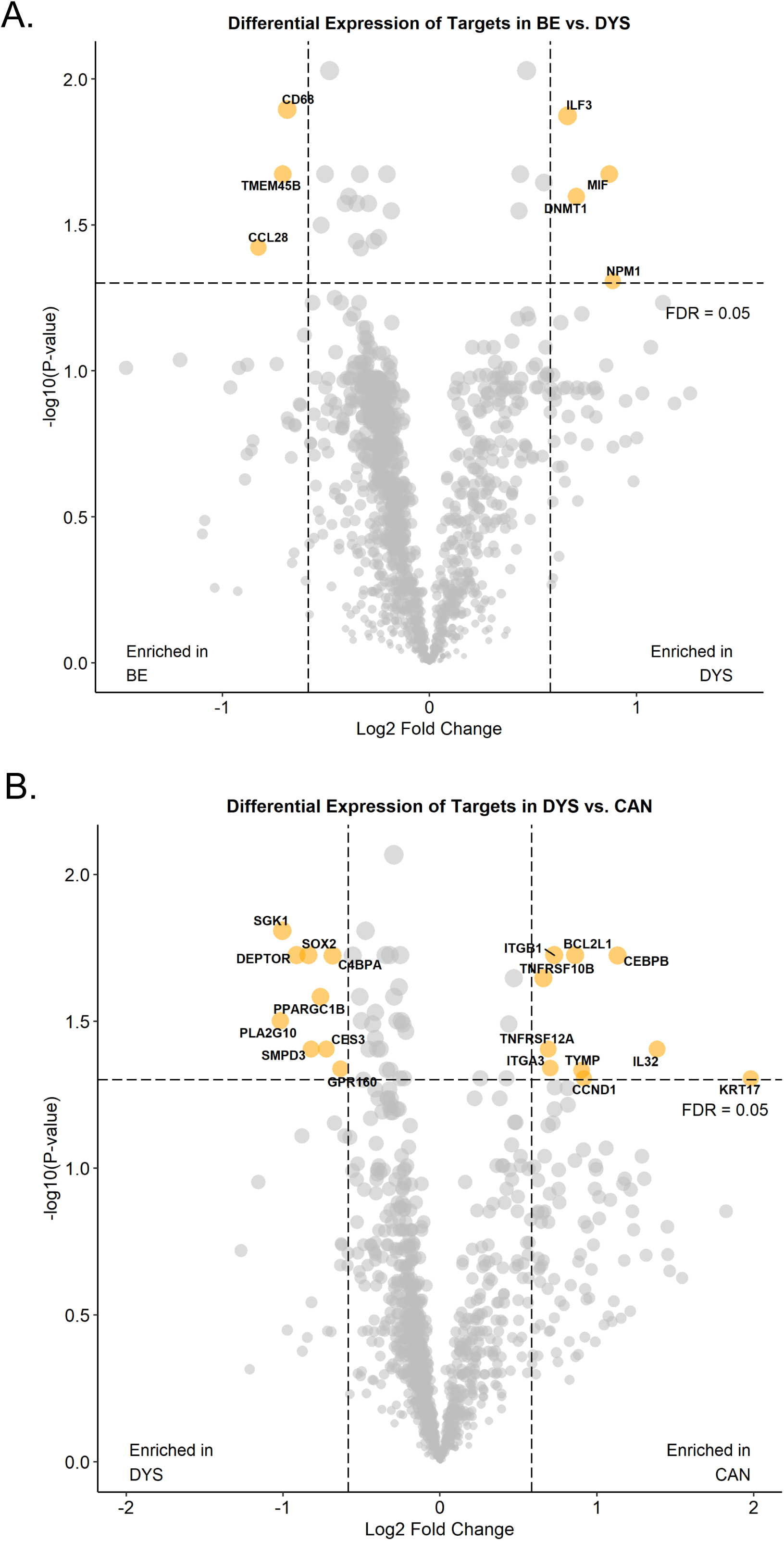
Differential RNA expression across stages of disease. Volcano plots showing differential RNA expression in non-dysplastic BE versus dysplastic samples (A) and dysplastic samples versus invasive EAC (B).

## Discussion

Characterization of the tumor microenvironment while maintaining spatial orientation has become an essential resource for the understanding of complex immune cell interactions and the assessment of biomarkers for prognosis and prediction of immunotherapy response. The Nanostring GeoMx DSP platform [19,20,25] allows for characterization of the tumor microenvironment through high-plex digital quantification of proteins and mRNA from FFPE tissues with spatial resolution [26]. The application of spatial technologies to large numbers of patient samples that are available in pathology archives as FFPE blocks potentially provides unparalleled insight into cell types, cell location, biomarkers, and the interactions that may underlie disease progression. This technology is rapidly growing and has been applied to cancer samples such as triple-negative breast cancer [27], lung cancer [28], and prostate cancer [29]. However, despite the success of spatial profiling within advanced tumors, pre-cancerous samples such NDBE have been understudied.

To explore the feasibility of studying NDBE to EAC progression through spatial profiling, we analyzed expressions of 49 proteins and 1800 RNA transcripts within small regions of the epithelium and stroma separately. Using this strategy, segmentation worked well based on differential expression and clustering analysis. As there is a trade-off between the number of cells profiled and signal, we benchmarked the performance of the protein panel and CTA utilizing areas typically coving ∼3 BE glandular profiles (Supplementary Figure 1E and 1F). While this approach was easy to do and provided a relatively large area for analysis, it did sacrifice some resolution.

For protein analysis, we demonstrated that the histone H3, GAPDH and S6 housekeeping probes correlated across ROIs, and were best used for normalization in the protein analysis. While IgG, surface area, and nuclei number were less reliable for normalizing data for quantification. Some previous studies have used area as a normalizer [30, 32]. However, area normalization does not take into account cellularity differences like housekeeping genes do. For our smaller biopsies that had variable cellularity, we found surface area and control antibodies to have relatively poor correlation compared to the housekeeping genes. Using the housekeeping genes for normalization, the protein assay seemed to provide results that would be expected and correlated with the transcriptomic analysis. Interestingly, using standard normalization and cutoff metrics for the transcriptome analysis suggested that the RNA profiling worked well with only 36/1,812 (2%) of transcripts considered below the LOQ in FFPE endoscopic mucosal resections. While the tissue pieces within these FFPE block are slightly larger than the endoscopic biopsies used for protein analysis, the ROIs that were actually analyzed were of a similar design and size.

Our pilot study utilizing NDBE biopsies from patients who would go on to progress to HGD/EAC and patients who did not progress suggested several interesting findings. CD66b (found on neutrophils) and CD68 (found on macrophages) were both increased in NDBE from progressors, suggesting myeloid cells may be important in the progression process. CTLA4, PD-1 and PD-L1 were also both increased in the stroma of progressor samples and CD45RO and HLA-DR were increased in the Barrett’s epithelium, suggesting that an immunosuppressive environment may develop early in the progression process. Given the limited number of samples utilized for this pilot, no strong conclusions can be made. However, these results suggest further studies confirming these findings, performing more detailed analysis on the subtypes of cells present, and determining if any differences may be useful as biomarkers for future progression are warranted.

In conclusion, this study highlights the utility of the NanoString GeoMax DSP for profiling relevant proteins and RNAs in small, FFPE endoscopic samples. Our pilot study of highly multiplexed measurements of protein expression suggested several myeloid and immune-modulatory markers were upregulated in biopsies from patients who would go on to progress to more advanced disease. The application of such novel platforms to provide comprehensive snapshots of clinical samples enables an unprecedented insight into the pre-cancer microenvironment that may be indicative of disease progression.

## Supplementary Materials

Supplementary Table 1 to this article can be found online.

## Author Contributions

Idea/Concept/Conceived of study (MDS, JB), Sample collection and annotation (NF, AMK, LCD, JB, LF), Histology review (RO, MDS), Performed data analysis (QA, MDS), critical review of data (MDS), wrote manuscript (QA, MDS), reviewed manuscript (NF, AMK, PS, RO, LF, LCD, JB). All authors have read and agreed to the published version of the manuscript.

## Acknowledgements

The authors acknowledge the NanoString team for performing the RNA and Protein analysis on the GeoMx DSP platform as well as assisting with data analysis as part of the TAP program and David Scoville (Nanostring) for critical review of the manuscript.

## Ethical Approval

All sample collection and studies were performed after IRB approval.

## Figure Legends

**Supplementary Figure 1:**
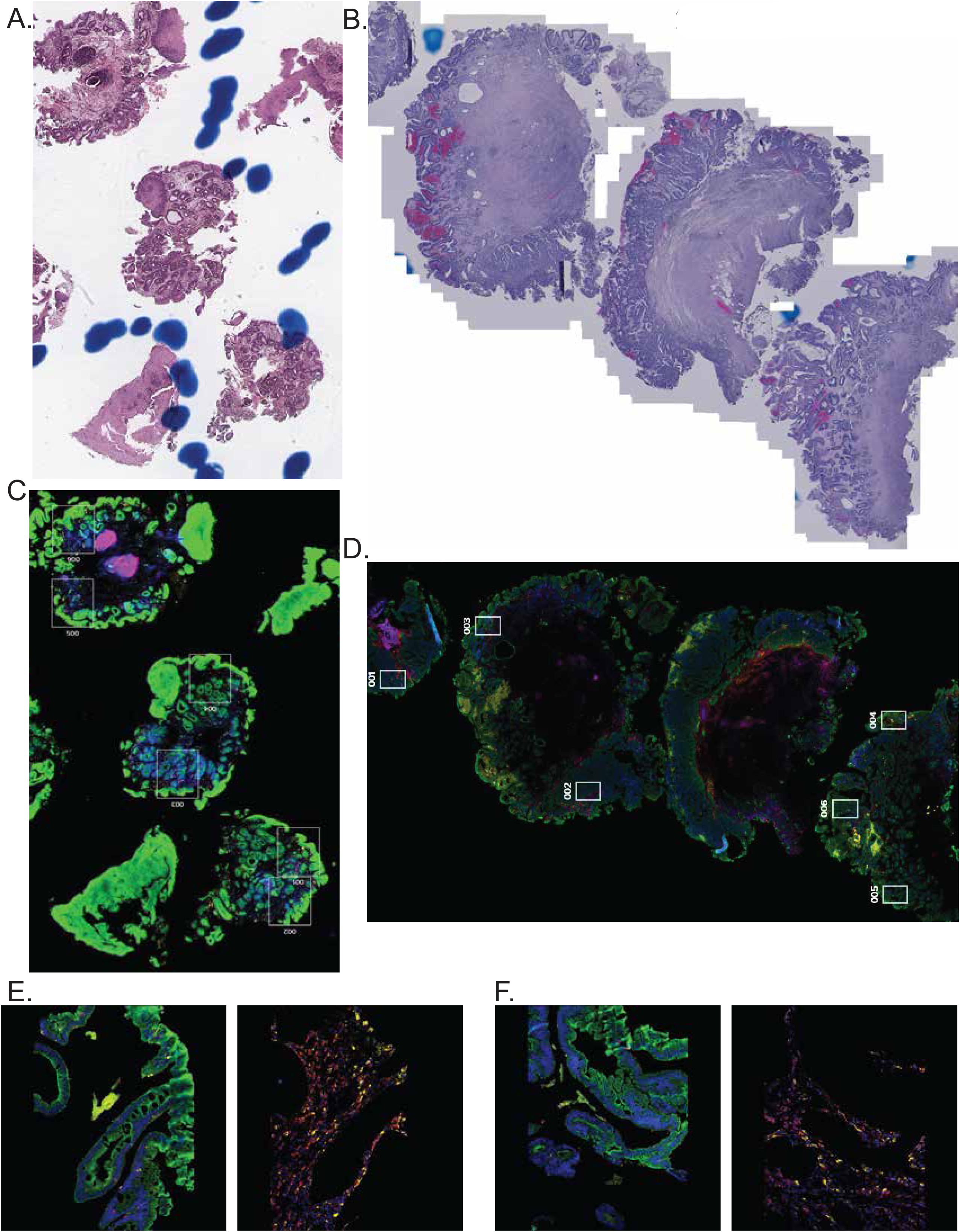
Representative images of BE biopsies and EMRs. Hematoxylin and eosin stained sections of endoscopic biopsies (A) and an EMR (B). Immunofluorescent imaging of the morphology markers used to select ROIs (A, biopsy) and (D, EMR). ROI segmentation into Pan-CK positive (left) and Pan-CK negative (right) AOIs in biopsies (E) and EMR (F). For C – F, Green: Pan-CK, Red: CD45, Yellow: CD68, and blue: nuclei.

**Supplementary Figure 2:**
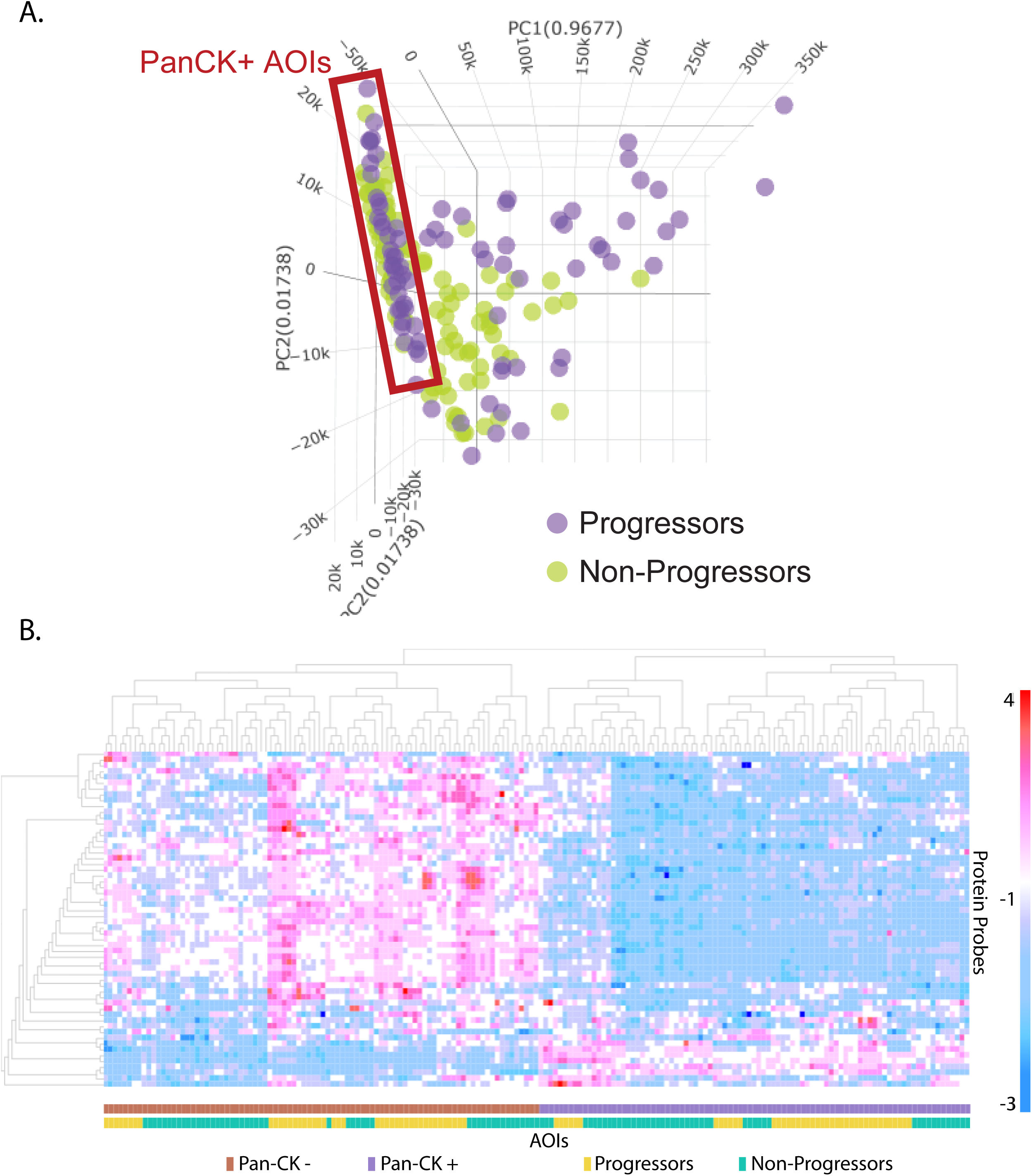
Segregation of AOIs in protein analysis. (A) Principal component analysis visualized in a three-dimensional tSNE plot. All Pan-CK+ AOIs tightly cluster (red box). AOIs from progressing patients and non-progressing patients do not strongly separate. Purple circles = all ROIs from progressive patients corresponding to group A. Green symbols = all ROIs from group B; non progressors patients. (B) Heatmap view of unsupervised clustering of protein AOIs. Pan-CK status perfectly segregated AOIs whereas progressing patients and non-progressing patients had less segregation in both pan-CK+ epithelial and pan-CK-stromal AOIs.

**Supplementary Figure 3:**
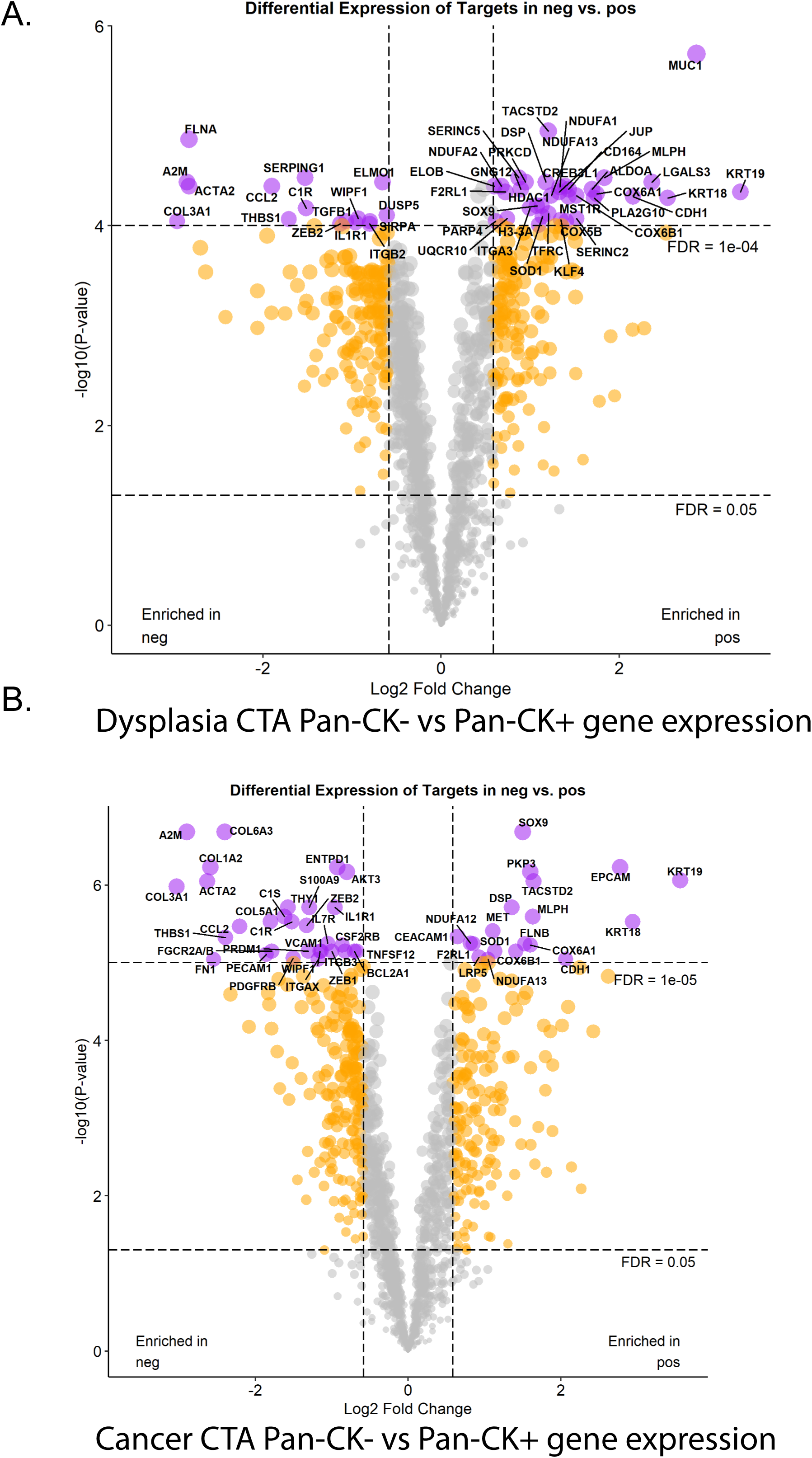
Segregation of AOIs in RNA analysis. Volcano plots showing differential gene expression in Pan-CK-versus Pan-CK+ AOIs in dysplasia samples (A) and EAC samples (B).

